## Supplementary materials for "Efficient Coding in the Economics of Human Brain Connectomics"

[1]Department of Neuroscience, Perelman School of Medicine, University of Pennsylvania, Philadelphia, PA 19104, USA
[2]Initiative for the Theoretical Sciences, Graduate Center, City University of New York, New York, NY 10016, USA
[3]Joseph Henry Laboratories of Physics, Princeton University, Princeton, NJ 08544, USA
[4]Department of Psychiatry, Perelman School of Medicine, University of Pennsylvania, Philadelphia, PA 19104, USA
[5]Department of Bioengineering, Schools of Engineering and Medicine, Stanford University, Stanford, CA 94305 USA
[6]Department of Psychology and Center for Brain Science, Harvard University, Cambridge, MA 02138, USA
[7]Penn-Children's Hospital of Philadelphia Lifespan Brain Institute, Philadelphia, PA 19104, USA
[8]Department of Neurology, Perelman School of Medicine, University of Pennsylvania, Philadelphia, PA 19104, USA
[9]Department of Physics & Astronomy, College of Arts and Sciences, University of Pennsylvania, Philadelphia, PA 19104, USA
[10]Department of Bioengineering, School of Engineering and Applied Sciences, University of Pennsylvania, Philadelphia, PA 19104, USA
[11]Department of Electrical & Systems Engineering, School of Engineering and Applied Sciences, University of Pennsylvania, Philadelphia, PA 19104, USA
[12]Santa Fe Institute, Santa Fe, NM 87501, USA
[†]Co-senior authors

September 13, 2021

### 1 Supplementary Modeling/Math Notes and Results

In this section, we investigate the use of a stochastic process to model the randomness and redundancy of brain network communication. Specifically, we assess the stochastic process of a random walk; in the network neuroscience literature, this process is also (somewhat confusingly) called the *diffusion model.* Adding to an extensive literature that models neurotransmission of signals as discrete stochastic events which propagate across neural circuits, we describe evidence from neuroimaging data that supports similar assumptions for macroscale brain network communication.

#### 1.1 Shortest path routing versus random walk models of network communication

In the subfields of computational and systems neuroscience, novel brain network communication models are needed. The key reason for this need lies in the fact that the predominant theory for network communication – a spectrum spanning from shortest path routing to random walks – has been criticized as both infeasible and inefficient [1, 2, 3]. In shortest path routing, neural signals travel from source to target using either the fewest connections or the shortest spatial distance [3]. Shortest path routing assumes that a single region has complete knowledge of the full network, allowing the region to choose to send messages along a shortest path, either topologically or physically. Clearly, this global knowledge is biologically infeasible. The infeasibility of the shortest path routing model's assumptions motivates the development of a model that does not depend upon this assumption. Random walk models are a promising alternative. They do not assume global omniscience, and instead propose a random propagation of information from source to target. Of course, it is true that random propagation leads to inefficiency. Yet, when viewed in the light of the biologically-established efficient coding framework, inefficiency can be powerful: the redundancy and stochasticity of communication by randomly walking messages are linked features that balance the transmission fidelity and lossy compression of information.

Despite its common criticisms, we test the shortest path model here in the supplement as an alternative to the random walk model evaluated in the main text. We do so because the shortest path model is commonly used as an explicit mathematical approximation of the communication efficiency of structural brain networks. Further, the diminutiveness of the average shortest path is thought to reflect the integrative capacity of a circuit [4, 5]. Intuitively, the shortest path between two regions is a direct connection. The presence of such a connection implies that those regions are sharing and integrating information. More recent models of shortest path communication move beyond these direct pairwise connections to multiple successive pairwise connections, and can thereby more comprehensively operationalize a network's integrative capacity. Such models have been called topologically optimal routing [1, 3]. A recent extension of topological routing models is called greedy routing, or navigability [6, 7, 8]. The navigability model assumes that a neural signal can triangulate its spatial, rather than topological, position to destinations in relation to the shortest spatial paths [6] or in relation to other network properties like degree and clustering [7]. In its favor, the navigability model does make fewer mathematical assumptions than an optimal topological routing model; yet, the biological support for those assumptions remains unclear. This navigability model is called *greedy* routing because it deterministically selects the optimal shortest route given local information in a hidden space [7, 8]. However, the hidden space is calculated based on global network properties. In contrast, the linear process of random walks – which we consider in the main text – is simple and decentralized, and has proven broadly useful in modeling diverse types of biological signaling [9, 10].

Shortest path routing models are commonly studied in network neuroscience as well as in systems neuroscience. In systems neuroscience, the circuit underpinning the computations serving a particular behavior is commonly defined as a subset of brain regions connected by shortest paths. Circuit schematics in the neuroscience literature implicitly test for and depict the direct (shortest) wiring diagram derived from biological tract tracing or similar techniques [11]. The weaknesses of shortest path routing become particularly apparent when we consider selectively associating a few localized pathways (out of tens of thousands of connections in the macroscale connectome and orders of magnitude more in the micro-scale connectome) with cognitive processes and resultant behavior. A more complete model that would address those weaknesses could consider the fact that the circuit is embedded in the full network [12], and thereby account for multi-hop transmission as it contributes to cognitive processes and behavior [13, 14].

In sum, in the network neuroscience literature, the shortest path routing model is made mathematically explicit, but there is scarce prior evidence of its biological relevance and questions regarding the biological plausibility of its assumptions [15]. We tested the shortest path model in our analyses with substantially greater statistical power than prior reports and found a lack of evidence for the model (Main Text Figure 2; Results 1.3) [1].

#### 1.2 Random walk dynamics as a model of network communication

The choice to model neurotransmission at the macroscale using random walkers is supported by models of signal diffusion along white matter fibers that can be measured through diffusion weighted imaging. The probability that a signal propagates along white matter fibers is directly proportional to the fiber's microstructural integrity. Prior work demonstrates that such models of signal diffusion can explain statistical dependencies among regional activity time series, as commonly summarized in measures of functional connectivity [16, 17]. Moreover, such models have been shown to explain how structures supporting random walks can predict functional connectivity [18], which we replicate in our own data in Supplementary Figure 4. Interestingly, the random walk model can also predict the direction along which neural signals flow [13]. Importantly, models of signal diffusion can also explain the trans-synaptic spread of pathogens in human and non-human connectomes [19, 20, 21, 22, 23, 24]; these studies provide important neurobiological validation for the use of random walk models in studies of macroscale brain organization and function.

The abstraction of a neural signal or message into a notional unit like a random walker is similar to the usage of quantal units in neuroscience. For instance, quantal units help us understand how discrete message packets of synaptic transmission of vesicles containing neurotransmitters encode patterns of information, such as quantal units of light [25, 9]. In our study, we do not venture to explain how neural coding occurs within brain regions to encode behaviorally relevant information. Rather, we ask the tractable question of how the brain network can reliably transmit messages in a simple repetition code [9]. Our focus on transmission allows us to circumvent the known challenges of using information theory to describe measurements of activity [26], by instead using the theory to posit and test a computational model of transmission. Specifically, we use the computational model of efficient coding – and more precisely, the repetition coding variant of redundancy reduction. This choice allows us to reframe the question of information processing as information transmission, which is important because the limits of computation and the limits of communication are intertwined [27].

Here we consider a communication model of diffusion, or random walks. A diffusion process can occur at multiple spatial levels and can be modeled in discrete- or continuous-time. Despite this diversity, what is shared by all diffusion processes is stochasticity [9, 28, 29, 30]. Here we model a stochastic process at the macroscale using a *random walk model* of discrete impulses. The discrete stochastic impulses are random walkers, which represent the combined contribution of signaling mechanisms including action potentials and non-neuronal processes [31, 32]. Below, we discuss biological interpretations of this model.

Perhaps the best understood diffusion process in the brain is the micro-scale biophysical diffusion of neurochemicals within cytoplasm and blood across short distances [9]. This kind of "analog" processing uses existing physical systems of the brain such as the random motion of neurotransmitters and neuromodulators interacting with channel receptors and cell structures. Such biophysical diffusion processes do not require neuronal materials and neuronal energy, which by contrast *are required* to transmit neural messages via action potentials along structural connections. Given this distinction in mechanism, we assess the role of micro-scale diffusion via blood in our analysis of biased random walks for CBF.

Across longer distances, "digital" processing requires the discrete pulses of action potentials to propagate neural messages. Electrical signaling depends on the voltage-gated channels that regenerate currents to propagate action potentials along axons. Towards this function, myelination of axons and the microstructural integrity of biological tracts directly impact signal conduction in white matter fibers and cortico-cortical communication [33, 34]. Therefore, we assess the role of longer-distance electrical signaling in our analysis of biased random walks for intracortical myelin content.

Together, the efficiency of metabolic diffusion (aided by CBF) and electrical signaling (aided by intracortical myelination) constitute the biological basis of our macroscale efficient coding model.

#### 1.3 Metabolic running costs of network communication architectures

We noticed that the effects of age on global efficiency and CBF were most drastic in childhood and early adolescence. This observation motivated us to perform a sensitivity analysis in which we split participants into two categories: those younger and those older than age 16. For participants younger than age 16 ($n$=555), we performed a Wilcox rank sum test to evaluate the relationship between the residual variance in CBF and global efficiency after regressing out age. We did not find a significant relationship between CBF and global efficiency ($W$=148330, $p$-value=0.29). For participants older than age 16 ($n$=479), we similarly did not find a relationship between the residual variance in CBF and global efficiency after regressing out age ($W$=122440, $p$=0.09). Our findings are distinct from those of others which reported a significant relationship between CBF and global efficiency when using a different group-splitting criteria [15]. Specifically, in that prior study, Group 1 consisted of students (n=11, ages=21-32 years, mean=25.4 years, SD=3.4 years, 5 females) and Group 2 consisted of a sample of adults (n=12, ages=23-57 years, average age=36.7 years, SD=10.7 years, 7 females; data acquired after a scanner upgrade). Collectively, our results indicate that age confounds the relationship between global efficiency and CBF, and the data does not support the claim that shortest path routing is associated with reduced metabolic expenditure.

#### 1.4 Adaptive trade-offs between metabolism and network architecture

Communication between brain regions or modules requires reliable transmission of information with an expected fidelity [27, 9, 3]. Although our data does not link metabolic expenditure to shortest path routing, communication of information diffusing along shorter paths should nevertheless confer advantages in speed and signal fidelity compared to longer paths. To test this hypothesis, we investigated whether brain metabolism is associated with network structures that support diffusion over shorter paths. Specifically, we assessed the association between CBF and path transitivity, a measure of the density of connections re-accessing shortest paths, thereby guiding diffusion along efficient pathways (Main Text Figure 5A; Supplementary Figure 2A (left)). Prior reports have demonstrated that path transitivity in structural networks is positively correlated with fMRI BOLD functional connectivity [18], a finding that we replicate in our own data (Supplementary Figure 4A). Path transitivity requires more connections and presumably incurs greater metabolic running costs associated with both the structural connections and increased functional connectivity [35]. When considering variation across individuals, we find that greater path transitivity is associated with greater CBF (Supplementary Figure 4B; $t$ = 2.27, estimated model $df$ = 11.45, $p$ = 0.02; controlling for age, sex, age-by-sex interaction, degree, density, and in-scanner motion). This result suggests that brain networks may strike a compromise between metabolic cost and the signaling advantages of path transitivity. Next, we sought to assess whether the relationship between brain metabolism and path transitivity was moderated by development. We found that the interaction between path transitivity and age was positively associated with CBF ($F$ = 24.6, estimated $df$ = 3.13, $p < 2 \times 10^{-6}$; Supplementary Figure 4B). Increased metabolic expenditure associated with greater path transitivity was prominent during adolescence, when global CBF tends to decrease [36].

We expanded our analysis of compromises between brain metabolism and network topology by considering multiple trade-offs. Specifically, we considered variations in metabolic cost, path transitivity, and modularity across individuals (Supplementary Figure 2A-B). We found that the relationship between path transitivity and global CBF is moderated by modularity ($t = 2.56$, estimated model $df = 13.43$, $p = 0.01$). When we consider variation across individuals, we find a saddle point function of metabolic costs, where the means for both path transitivity and modularity fall at the saddle point (Supplementary Figure 2C). A saddle point suggests that adaptive compromises in network architecture are constrained by dual objectives. Along one axis, the objective is minimizing metabolic expenditure by coupling modularity with path transitivity. Along the other axis, the objective is maximizing metabolic expenditure by decoupling modularity from path transitivity. Brain networks may negotiate multiple trade-offs between metabolism and structure such that most brain networks reside around a saddle point with locally optimal metabolic savings when network structure is coupled, whereas a smaller fraction of brain networks reside at a global minimum with globally optimal metabolic savings when network structure is decoupled. An analysis of the landscape relating CBF, modularity, and path transitivity for brain regions suggests that the clustering of brain regions that contributes to modules is associated with greater metabolic cost, whereas the clustering of brain regions neighboring shortest paths is associated with reduced metabolic cost (Supplementary Figure 2D). Random walk models of network communication explain how structures that prioritize the probability of taking certain pathways (like shortest paths) can predict functional connectivity [18], and we replicate this finding in our own data (see Supplementary Figure 4).

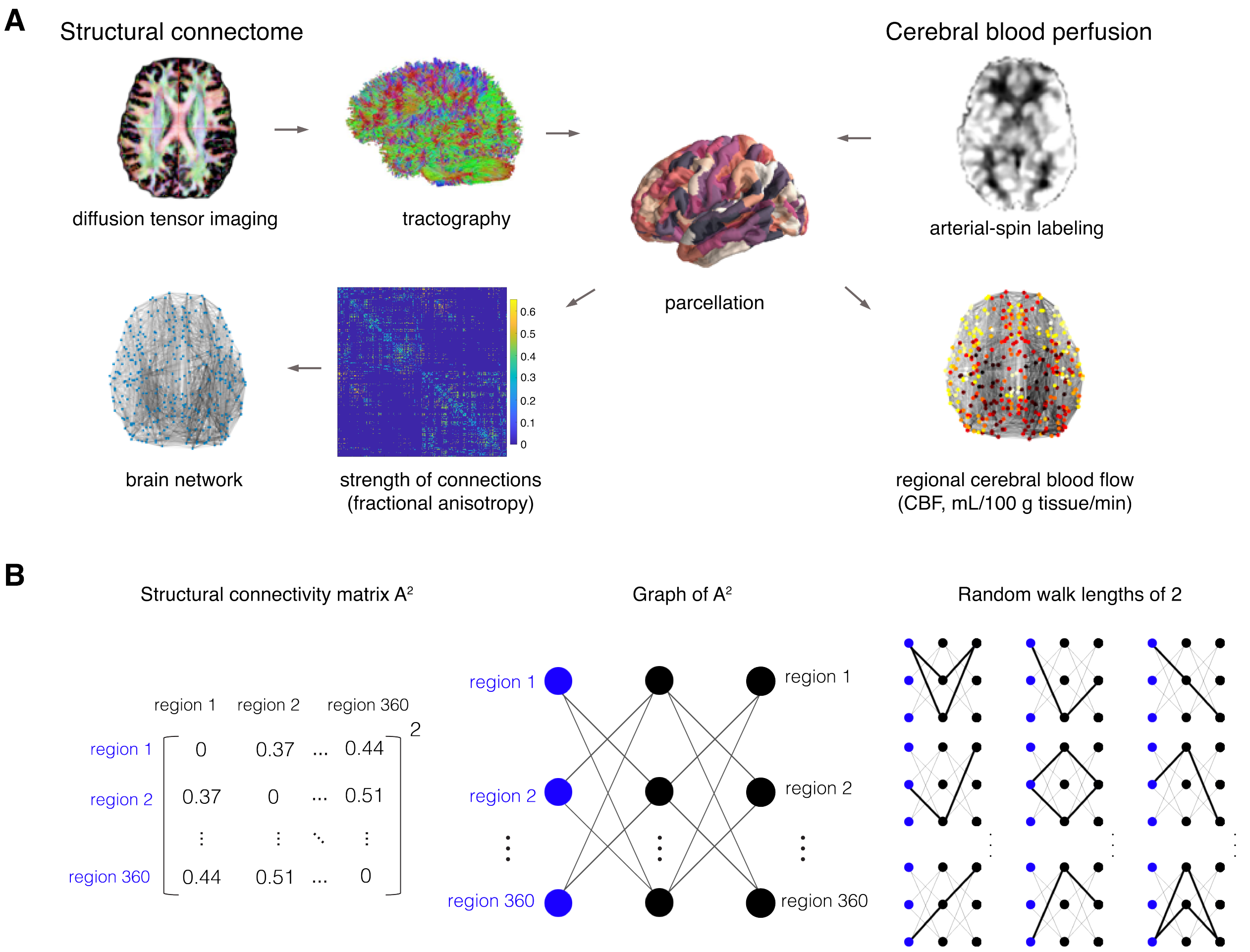


Figure 1: **Schematic overview. (A)** Image acquisition and processing pipeline, generating structural connectomes using deterministic tractography of diffusion spectrum imaging data and extraction of cortical regions using the Glasser anatomical atlas. Regional cerebral blood flow (CBF) during a resting state was quantified using arterial spin labeled perfusion MRI at 3T. **(B)** Structural networks are represented by a connectivity matrix (left). Each row and column (node) designate distinct brain regions. Each cell (edge) indicates the integrity of the white matter connection, measured with fractional anisotropy. In matrix multiplication, the elements of the calculation can be represented graphically by the row and column vector elements as edges (middle). For example, the matrix to the exponent of 2 reflects the overall integrity of structural connections spanning a random walk length across two connections (right). The solution is the dot product, multiplying the edges comprising each possible path of length 2 and summing the resulting value(s). Intuitively, in the context of brain structural connectomes, the resulting matrix values represent the integrity of connections composing paths with a random walk length equal to the exponent's integer value.

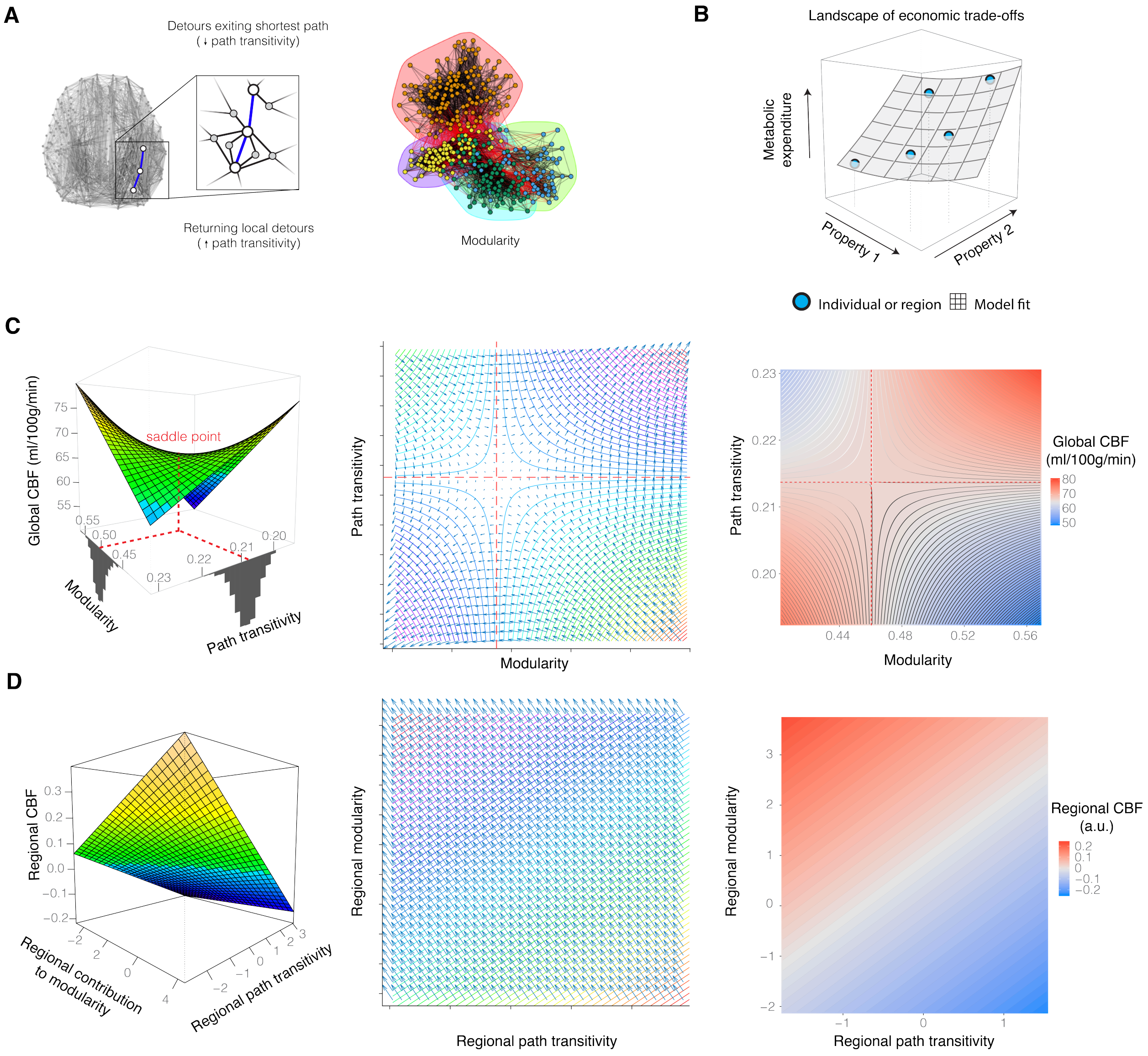


Figure 2: **Metabolic savings associated with regional connections contributing to integration and lossy compression.** **(A)** Greater path transitivity enhances the ability of diffusing signals to re-access the shortest path when the number of closed paths (triangles) returning to nodes on the path is high. Path transitivity of the brain's structural network is statistically associated with fMRI BOLD activity [18]. Modularity is a feature of brain organization across species whereby regions cluster into highly intraconnected communities. Modularity is thought to confer efficient use of physical materials, segregate information representation, and support efficient information transfer among brain regions. **(B)** The 3-dimensional fitness landscape conveys adaptive trade-offs associated with brain metabolism in the context of evolutionary constraints on efficiency. **(C)** When we consider variation across individuals, we find that economical network architectures are adaptively balanced with metabolism. Metabolic cost is associated with the interaction of path transitivity and modularity ($t = 2.56$, estimated model $df = 13.43$, $p = 0.01$), controlling for age, sex, age-by-sex interaction, degree, density, and in-scanner motion. The mean path transitivity and modularity across all individuals (red dotted lines projected from frequency histograms) approach a saddle point, defined as a point on the surface that is both a relative minimum and a maximum along different axes. We calculated a map of change in global CBF with respect to both modularity and path transitivity using first-order derivatives. To quantify the location of the saddle point coordinate within the change map, we performed a $k$-nearest neighbor search of the value 0 in the gradient of first-order derivatives indicating minima and maxima (see Figure 2). The saddle point suggests that adaptive compromises in network architecture are constrained by dual objectives. Along one axis, the objective is minimizing metabolic expenditure by coupling modularity with path transitivity. Along the other axis, the objective is maximizing metabolic expenditure by decoupling modularity from path transitivity. This landscape suggests the existence of compromises which balance adaptations of functional flexibility and spatial efficiency with material and metabolic costs. **(D)** Regions with greatest metabolic expenditure tend to have reduced

Figure 2: **continued.** average path transitivity ($t = -2.76$, $p = 0.006$). The regional contribution to modularity and the average regional path transitivity explained differences in regional CBF ($F = 6.2$, $R^2$=0.03, $p = 0.002$, $df = 357$).

#### 1.5 Metabolic expenditure associated with rich-club hubs

If *functional* hubs demand greater metabolic expenditure [35] and functional connectivity explains a portion of the variance in structural connectivity [37, 38], then a reasonable hypothesis is that structural hubs also demand greater metabolic expenditure. However, there is sparse evidence to support this hypothesis [39]. To test the hypothesis that structural rich-club hubs demand greater metabolic expenditure, one previous study performed an analysis testing for a trend between regional structural connectivity of hubs and metabolic cost [39]. The investigators employed an exploratory comparison of (1) an atlas of metabolic expenditure created from one dataset and (2) structural connectivity of a sample of adults from a second dataset (n=42, mean age= 29 years, SD=8 years, 16 females). The authors used the mean metabolic expenditure of each region in the atlas to assign a metabolic expenditure to rich-club hub and other brain regions of a separate group of individuals. With this method, the investigators found that rich-club nodes had greater metabolic expenditure than non-rich-club nodes.

We built upon this finding and re-assessed whether rich-club hubs have greater metabolic expenditure than non-rich-club hubs across youth. Rather than using an atlas defined from separate individuals, we assessed metabolic and structural connectivity data from the same individuals by operationalizing metabolic expenditure with CBF (n=1,042). In contrast to the existing exploratory analysis, we found some evidence of *reduced* CBF in rich-club hub versus non-hub nodes (Supplementary Figure 8A). However, after controlling for age, sex, and in-scanner motion, we did not find evidence that rich-club nodes had greater metabolic expenditure than non-rich-club nodes (Supplementary Figure 8B). Our analysis was better statistically powered to find a relationship, and our null finding warrants alternative explanations that reinterpret subsequent work studying the metabolic properties of rich-club hubs. Hubs develop very early [40], yet high energetic costs of neurodevelopment have typically been hypothesized to result in slow and protracted growth [41]. We assess these alternative explanations in light of the existing literature on the development of hub structure.

In our own data, we performed exploratory analyses to assess explanations consistent with the alternative hypothesis of reduced metabolic cost in structural hubs. We found results consistent with structural hubs having reduced metabolic cost due to having greater cortical myelin content. Using previously published maps of intracortical myelin, we found that the hubs of the rich club (colored in red in Supplementary Figure 8) which were stronger tended to have greater cortical myelin content (Spearman's $\rho$=0.35, $p$=0.02; Supplementary Figure 8B). In support of the notion that intracortical myelin of hubs contributes to their metabolic efficiency, we found that the hubs with greater cortical myelin content tended to have reduced metabolic expenditure (Spearman's $\rho$= -0.34, $p$=0.03; Supplementary Figure 8C). Together, greater cortical myelin and reduced CBF may explain why we found some evidence of hubs having reduced CBF compared to non-hubs (Supplementary Figure 8A). Given the established importance of neurodevelopment for cortical myelin levels [42, 43], we further evaluated our effects as a function of development. We found that the reduction of CBF in hubs versus non-hubs becomes statistically insignificant when controlling for age and sex (Supplementary Figure 8B).

In sum, the hypothesis that rich-club hubs have greater metabolic expenditure than non-hubs was not supported by our data. This null result suggests that at least one of the assumptions that originally motivated the hypothesis was spurious or requires more precision [35, 37, 38]. For instance, functional connectivity explains only a portion of the variance in structural connectivity [37, 38]. Therefore, we may not be justified in expecting that a prior report of a relationship between metabolic cost and functional connectivity [35] would carry over neatly to structural connectivity. The relationship between functional and structural connectivity in general is studied in work on structure-function coupling, where there is a general positive correlation at the whole-brain level [37, 38]. However, the relationship differs across regions, being weakest in frontal areas

[38]. Rich-club hubs are a particular class of highly connected regions in which structure and function are expected to be positively correlated. To corroborate this expectation in our dataset, we tested for a relationship between structural connectivity strength and BOLD functional connectivity strength (Supplementary Figure 8A). Consistent with prior work [38], we found that regions with greater structural strength tended to have greater functional strength (Spearman's rank correlation $\rho$=0.17, $p$ <0.001). From these data, we see that function explained about 2.9% of variance in structure. Hence, one explanation for our null result is that the complexity and heterogeneity of structure-function relationships thwart a key assumption that motivates the hypothesis that hubs have greater metabolic expenditure.

Inferences based on the assumption of a strong positive correlation between structural and functional connectivity are also complicated by previously studied brain network communication models [18]. The so-called diffusion models of communication in network neuroscience posit stochastic linear dynamics atop structural connectivity [44]. This model successfully explains variance in functional connectivity [18]. In contrast to the assumption of a positive correlation between structural and functional connectivity, this model predicts (and their data showed) that high-degree nodes (greater structural connectivity) attenuate the strength of functional connectivity because stochastic impulses have a higher probability of diverging from the shortest paths which contribute to functional activity [18]. Moreover, communication dynamics atop rich-club structure promotes low energy transitions in the control of functional network dynamics [45]. Using similar reasoning about the probability of propagating along versus diverging from shortest paths, we also found that hubs, due to their greater out-degree, require more random walkers to reliably send messages as a source node to other brain regions by the shortest paths (Main Text Figure 6B). Moreover, this prior study found that high path transitivity was associated with greater functional connectivity [18], which we replicated in our own data (see Supplementary Figure 4A). Path transitivity is a metric of transitivity of nodes comprising a shortest path. Prior theoretical work and our own analysis showed that hub nodes have the lowest transitivity in a hierarchically organized network [46]. In light of these findings which suggest that high degree attenuates the probability of reliable communication for a single stochastic impulse, that low path transitivity is associated with low functional activity, and that structural hubs have the lowest regional transitivity, it is unlikely that hubs have high functional activity given a diffusion-based communication process.

Another explanation for our null finding that rich-club hub CBF did not differ from non-hubs comes from prior literature on the neurobiology and neurodevelopment of intracortical myelin [47, 42]. In development, hubs have been reported to undergo the greatest rate of cortical thinning and myelination [42, 43]. Notably, however, prior analyses of baseline myelination of hubs versus non-hubs at age 14 years did not find evidence of a relationship between myelination and regional degree (see reference [42]'s Supplementary Figure 4). In our own data, we found that CBF was not correlated with node degree ($p_{\mathrm{SPIN}} > 0.05$). These null results are consistent with the null hypothesis that rich-club hubs do not differ in myelination from non-hubs and thus do not differ in metabolic expenditure either.

The last explanation we explore proposes the alternative hypothesis that rich-club hubs with long-distance connectivity have reduced metabolic expenditure compared to non-hubs because myelination supports metabolically efficient neural communication. We explored the explanation that long-distance connections (like those in the rich-club hubs) tend to have greater myelination than other brain regions, which would support greater metabolic efficiency [48, 1, 49, 50, 51, 42]. In our data, the structural connections of the rich-club hubs had greater average length than that of non-hubs ($t = 138.7$, $df = 2080$, $p < 0.001$). Myelin is more prevalent in white matter than gray matter [48, 50, 51], and accordingly white matter expends less metabolic resources than gray matter for signaling [52]. Following the results of a prior report [50] and the notion that FA is a microstructural index for physical properties of white matter [53, 54, 55, 56, 57], we used our own data to replicate a negative correlation between CBF and the strength of a node's structural connectivity ($r = -0.15$, $p = 0.003$, $df = 358$). Together, rich-club hubs may have reduced metabolic expenditure because they have connections of greater average length and FA compared to non-hubs, which prior work suggests also have greater myelination.

##### 1.6 Cognitive efficiency, speed, and accuracy

The compression efficiency of rich-club hubs, and any subset of brain regions, depends on how the connectivity of the whole network shapes random walk dynamics. Hence, cognitive efficiency being associated with compression efficiency in other non-hub brain regions is consistent with the importance of diverse dynamics generated by the whole network [58, 59]. In support of the importance of whole network connectivity, we found that individuals with reduced compression efficiency, prioritizing transmission fidelity, of brain regions outside the rich club tend to exhibit increased cognitive efficiency of complex reasoning ($t = -4.72$, bootstrap 95% CI [-6.63, -2.72], model adjusted $R^2$=0.21, bootstrap 95% CI [0.16, 0.26], estimated model $df = 10.50$, $p = 2 \times 10^{-5}$), executive function ($t = -2.85$, bootstrap 95% CI [-4.85, -1.0], model adjusted $R^2$=0.50, bootstrap 95% CI [0.46, 0.56], estimated model $df = 10.64$, $p = 0.03$), and social cognition ($t = -2.30$, bootstrap 95% CI [-4.21, -0.33], model adjusted $R^2$=0.22, bootstrap 95% CI [0.17, 0.26], estimated model $df = 10.45$, $p = 0.04$). Connectomes prioritizing fidelity tended to perform with greater cognitive efficiency in a diverse range of functions [60].

Lastly, we sought to investigate performance accuracy and speed separately. The theoretical model predicts that distinct properties of the connectome support either transmission fidelity or lossy compression, manifesting as greater accuracy or speed of behavioral performance. To test this prediction, we evaluated the more precise hypothesis that the compression efficiency of the network defines the minimum required information to be transmitted for a given fidelity, and should thereby correlate with accuracy. Moreover, shortest path complexity (path transitivity) may support a low-fidelity regime, and should thereby correlate with speed. To assess these predictions, we used a prior factor analyses of the cognitive battery of 14 tasks with latent variables separately modeling accuracy and speed. This factor analysis supported a 3-factor solution for accuracy and 3-factor solution for speed [61]. The 3-factor accuracy variables corresponded to executive and complex cognition, social cognition, and memory. The 3-factor speed variables corresponded to fast speed (e.g., working memory and attention tasks requiring constant vigilance), episodic memory speed, and slow speed (e.g., tasks requiring complex reasoning). We found that the accuracy of performance in executive and complex cognition was negatively associated with compression efficiency ($t = -4.58$, bootstrap 95% CI [-6.61, -2.52], $df = 5$, $p < 0.001$, $R^2$=0.041, bootstrap 95% CI [0.021, 0.075]; **Figure 9C**), whereas the speed of performance on the fast-speed tasks was negatively associated with shortest path complexity ($t = -2.12$, bootstrap 95% CI [-4.17, -0.05], $df = 5$, $p = 0.03$, $R^2 = 0.019$, bootstrap 95% CI [0.01, 0.044]; **Figure 9D**). We did not find relationships between compression efficiency and the other separated speed and accuracy factors (see **Supplementary Figure 8**). Taken together, networks with compression efficiency prioritizing fidelity tended to exhibit greater accuracy in executive and complex cognition, while networks with reduced shortest path complexity prioritizing lossy transmission tended to exhibit greater speed in fast-speed tasks.

#### 2 Supplementary Discussion

Brain systems viewed as information processors exhibit recurring compromises between information efficiency and other resource costs at the cellular [69, 70] and circuit levels of the brain [71, 72]. At the neuronal level, an optimal strategy for distributed coding is to reduce population size while distributing activity among a fraction of cells [69, 70]. Brain networks may reach a similar compromise through information processing constraints on complexity (i.e. size) of the network and its modules, and increasing the number of endogenously active components, such as in the default-mode system. Efficient coding predicts that bit rate varies as a function of the number and redundancy of synapses [70, 73]. Transmitting the same message across many parallel paths improves fidelity and increases bit rates, but information rate increases sublinearly with the number of paths because the system is highly redundant, incurring greater metabolic costs [70]. We similarly found that individuals with greater path transitivity—more redundant and lossy alternative paths to the shortest, direct paths—tended to require sublinearly increasing computational costs and tended to

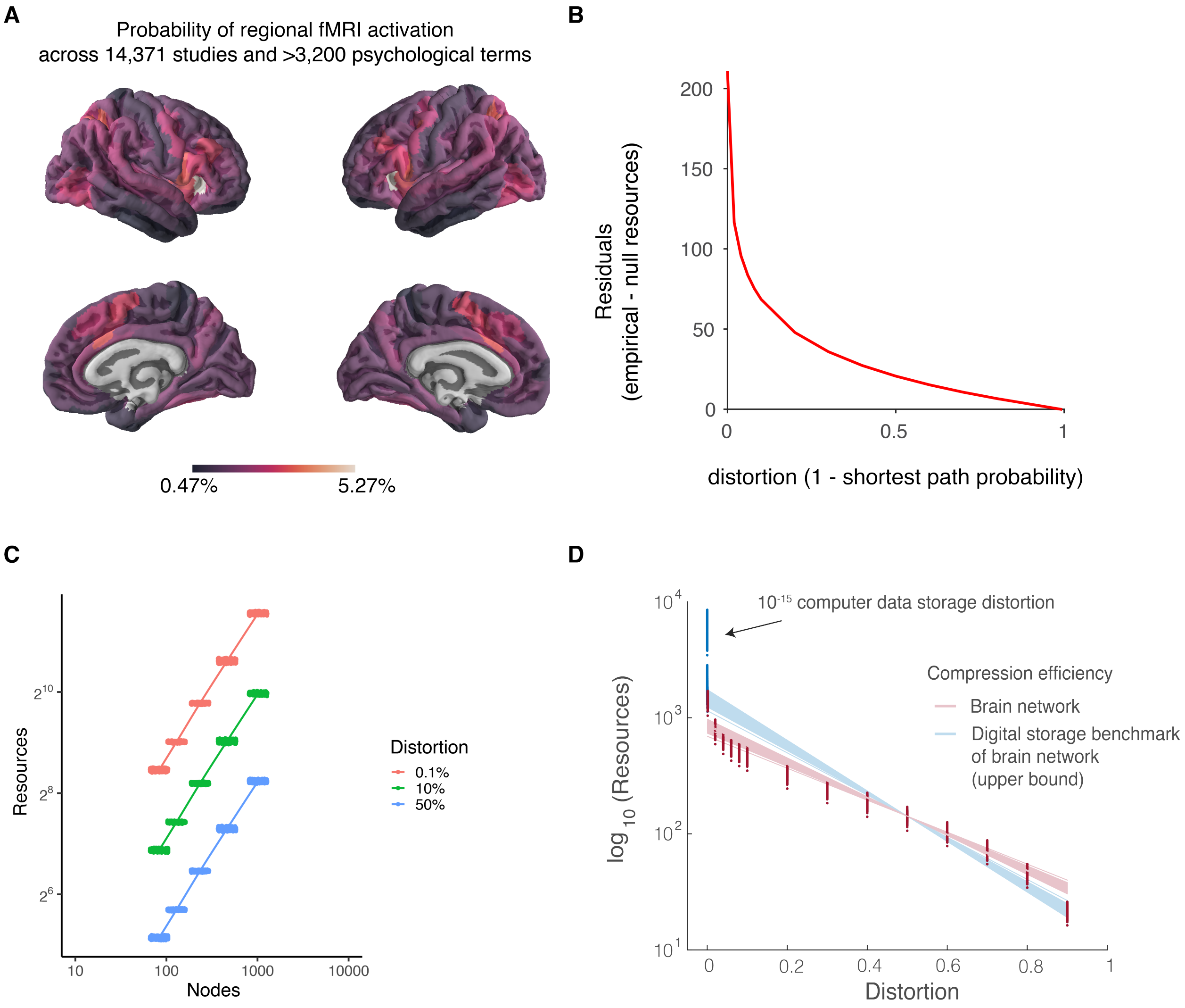


Figure 3: **Information processing in brain networks.** **(A)** Efficient coding posits communication of compact representations of information, which we model using the theory of lossy compression. To measure the information content of a neural message, which we model with a random walker, we require an estimate of the probability of a communication event. Forward inference probabilities characterize the chance that a brain region is active conditional on a particular set of behavioral or cognitive functions. Forward inference maps were automatically generated during the NeuroSynth meta-analysis (Main Text Method 7.5.6; Equation 16). Specifically, we obtained the probability of observing activation given the presence of a psychological term *P(Activation|Term)* across 14,371 studies and over 3,200 psychological terms [62]. Here, we use the average regional probability to calculate the information content carried by a random walker. The main results of the paper do not depend on the specific values estimated of information content, because each random walker conveys the same information content on average, so the number of random walkers will be scaled by this same value. Hence, we use simple independence assumptions to produce the estimate of average information content. Future work could assess different priors over psychological terms to refine the theoretical estimates of information content in accordance with empirical measurements in fMRI data. **(B)** A residual plot of Figure 3D in the main text. As distortion decreases, the empirical brain networks require exponentially greater resources than Erdös-Renyí random networks. **(C)** Information processing costs and capacity grow monotonically with random network complexity. Random networks also exhibit monotonically increasing resource costs as a function of network complexity, which we operationalize here as network size. As networks grow larger, their information processing costs and capacity grow in tandem. **(D)** Brain network information transfer does not achieve the design benchmarks of digital storage devices. To demarcate an upper bound of compression efficiency, we solved for the minimum resources required to achieve $10^{-15}$ distortion, a benchmark for digital storage devices [63]. The

Figure 3: **continued.** distance between the predicted rate-distortion gradient and the observed minimum resources required for the digital benchmark in brain networks suggests that the digital benchmark is impractical for brain networks. Instead, the rate-distortion gradient suggests that our selected upper bound of $10^{-5}$ distortion is a reasonable analysis choice for brain networks. We sought to establish an upper bound for the rate-distortion gradient. An upper bound prevents the slope from arbitrarily increasing depending on the resources corresponding to our choice of the minimum level of distortion. To do so, we compared the resource efficiency gradient in brain networks against a benchmark distortion level of $10^{-13}$ percent for digital storage devices [63]. Regional compression efficiency can be computed from two perspectives: brain regions are message senders or message receivers. For both senders and receivers, resource costs deviated from the rate-distortion gradient at extremely low distortion levels from 0.1 to $10^{-13}$%, suggesting that a reasonable rate-distortion gradient upper bound in brain networks is 0.1% distortion.

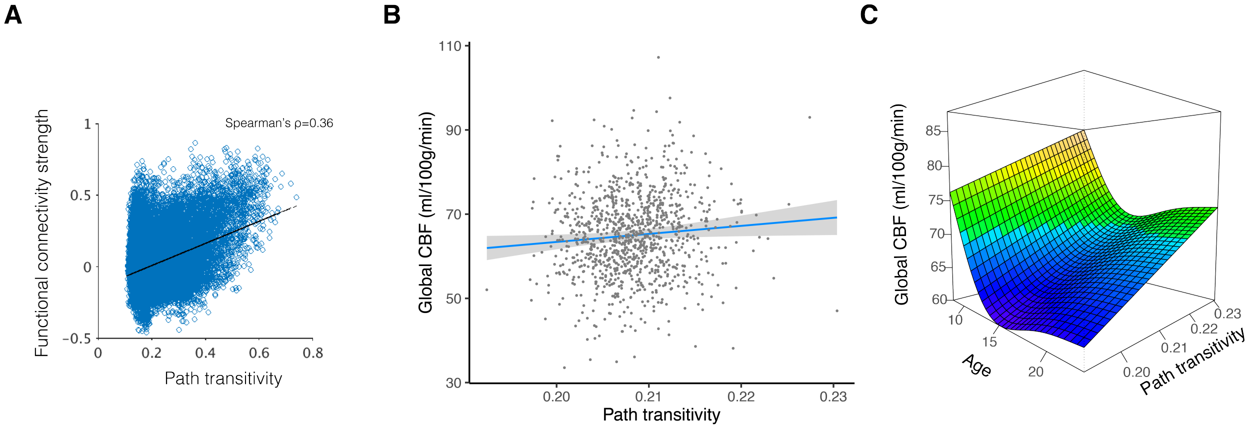


Figure 4: **Path transitivity is correlated with task-unconstrained functional connectivity strength and CBF. (A)** Here we replicate a previous report that path transitivity predicts functional connectivity strength [18]. Functional connectivity was measured using BOLD fMRI in a task-unconstrained state. Path transitivity and connection strengths were averaged across all participants. **(B)** When considering variation across individuals, we find that greater path transitivity is associated with greater CBF ($t = 2.27$, estimated model $df = 11.45$, $p = 0.02$), controlling for age, sex, age-by-sex interaction, degree, density, and in-scanner motion. Brain networks may strike a compromise between metabolic cost and the signaling advantages of path transitivity. **(C)** The interaction between path transitivity and age was positively associated with CBF ($F = 24.6$, estimated $df = 3.13$, $p < 2 \times 10^{-6}$; Supplementary Figure 4C). Increased metabolic expenditure associated with greater path transitivity was prominent during adolescence, when global CBF tends to decrease [36].

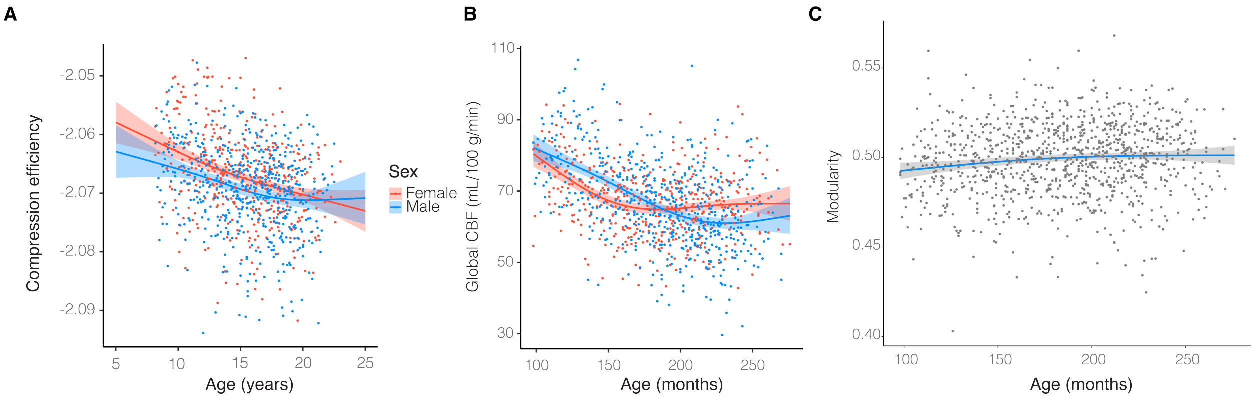


Figure 5: **Neurodevelopment places a premium on fidelity. (A)** When arbitrarily including increasingly lower levels of distortion to 0.001%, compression efficiency values increase due to the exponentially greater resource cost of reducing distortion. However, we find largely converging results as with our selection of a minimum distortion of 0.1%. Here, we show the finding of Figure 3F in the main text, that neurodevelopment prioritizes communication fidelity. **(B)** Global CBF evolves in a non-linear U-shaped trajectory with development. Here we replicate a previous report of the effect of age and sex on the evolution of global CBF with development [36]. CBF in females declines more rapidly in childhood than CBF in males. In adolescence and adulthood, global CBF is greater in females than in males. Males exhibit a decline in global CBF through childhood and adolescence, which plateaus into adulthood at a level lower than that of females. **(C)** Modularity increases with development. Here we replicate a previous report of the effect of age on modularity. Modularity tends to increase with development [64].

have greater global metabolic expenditure (Supplementary Figure 4). Taken together, the efficient coding model addresses the notable absence of biologically plausible and efficient inter-regional brain network communication models [1, 3].

#### 2.1 Communication models and network structure

Current work in network neuroscience views the shortest path and diffusion models as opposite extremes of a spectrum of communication processes [44, 74, 3]. In practice, the result of such processes is inextricably linked with the network structure upon which they occur. To understand this link, it is often useful to study the role of particular network features. Such a study can be made more concrete by first clarifying a conceptual distinction. Specifically, we distinguish between the theoretical model of communication and the descriptive measures used to characterize the structure that might help implement that communication.
Our findings motivate future studies of information integration in connectomes that model information transmission as a random walk rather than shortest path routing [75, 76, 2, 3, 77, 78]. Connectome architecture and biology optimize the objective of repetition codes by minimizing redundancy. Towards this objective, repetition coding is constrained by developmental cost of network architecture and biology, balanced by the efficiency of redundancy reduction. Artificial networks with large probabilities of shortest path communication – such as Erdős-Renyí networks – can achieve larger redundancy reduction than the human connectome [44]. However, such random networks are biologically implausible due to high developmental and evolutionary costs [44]. Nevertheless, the human connectome's biological properties can enhance signal propagation towards the efficiency of this random network with a repetition code.

#### 2.2 Connection between communication models and meso-scale connectome architecture

Meso-scale structures of brain networks, such as small worldness and the rich club, each have a distinct influence on the descriptive measures of network architecture [79, 67, 44, 74]. Moreover, the precise values of

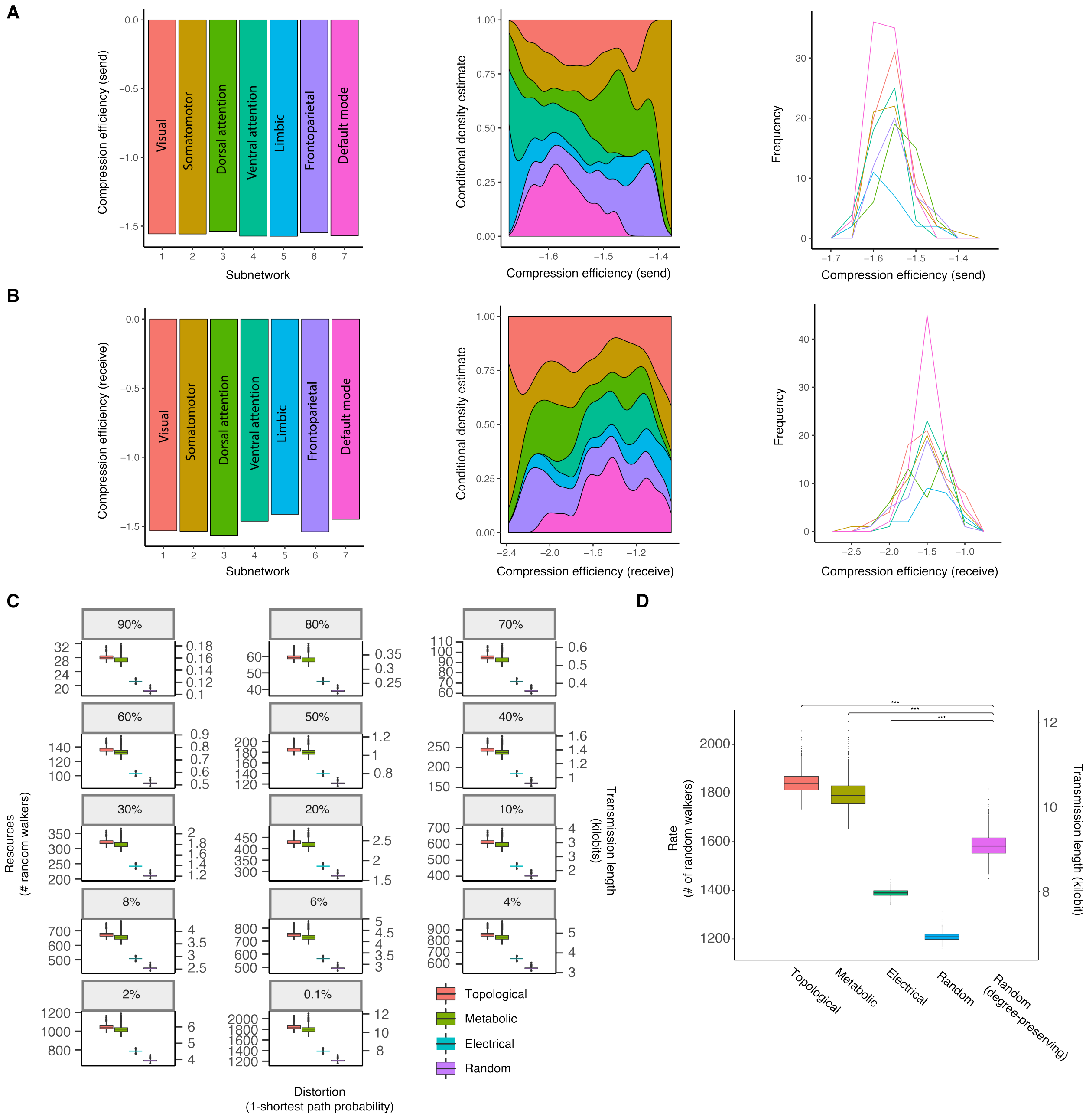


Figure 6: **Compression efficiency varies by subnetwork and connectome biology. (A)** Sender compression efficiency

Figure 6: **continued.** across subnetworks. Histograms depict the mean values (left). The conditional density estimates (middle) provide the probability of subnetwork values across the range of compression efficiency values. At high send compression efficiency, it is dominated by the somatomotor network. At lower send compression efficiency prioritizing fidelity, the default mode, frontoparietal, and attention networks appear dominant. The corresponding frequency of subnetwork send compression efficiency values are shown which were used to calculate the conditional density estimate (right). **(B)** Receiver compression efficiency across subnetworks. Histograms depict the mean values (left). The conditional density estimates (middle) provide the probability of subnetwork values across the range of compression efficiency values. At high receiver compression efficiency, the visual, somatomotor, and limbic networks represent a majority of values. At lower receiver compression efficiency prioritizing fidelity, the somatomotor and visual networks appear dominant. The corresponding frequency of subnetwork receiver compression efficiency values are shown which were used to calculate the conditional density estimate (right). **(C)** The finding reported in Main Figure 4 for 0.1% distortion generalizes across levels of distortion. **(D)** Compared to rewired null networks preserving the degree sequence, structural topology and metabolic resources support communication that prioritizes fidelity, while myelination supports communication that prioritizes compression. Asterisks denote $p < 0.001$).

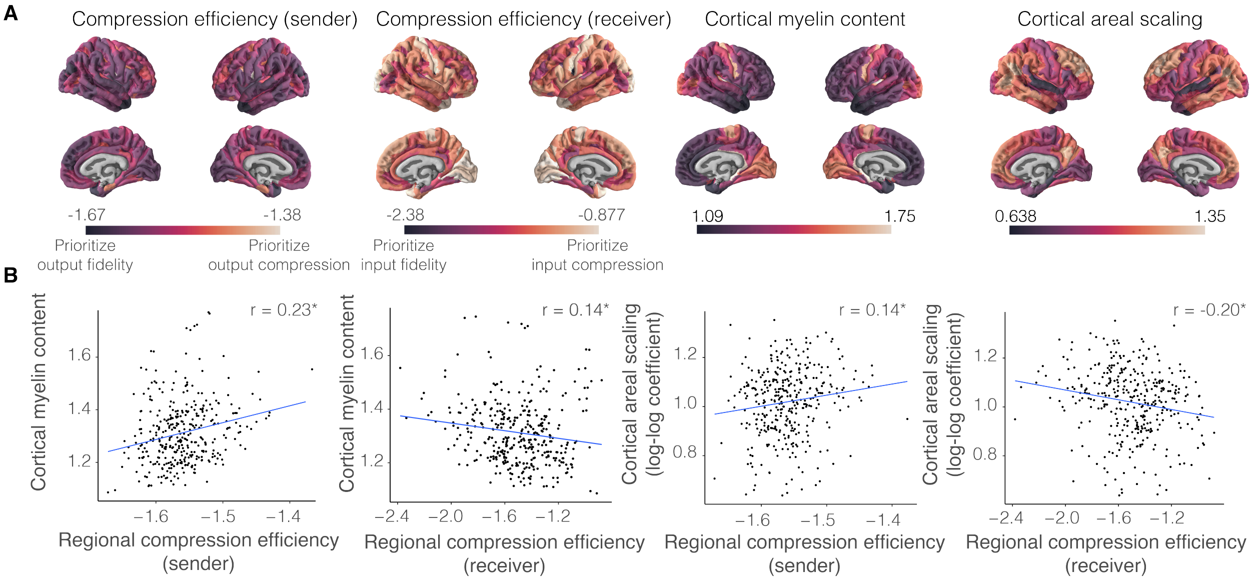


Figure 7: **Compression efficiency correlates with cortical myelination and areal scaling.** **(A)** Sender compression efficiency differs regionally across the cortex and describes the number of diffusing messages required to transmit information with specified signal fidelity (Supplementary Figure 6). Receiver compression efficiency describes the number of messages required to receive information with an expected signal fidelity. Regional values were averaged across individuals. **(B)** Brain regions differ in myelination and non-linear spatial scaling ratios of surface area change to total brain size change over development. Regional values were obtained from published maps [51, 65]. Cortical regions with greater levels of myelin tend to have greater sender compression efficiency ($r = 0.23$, $p_{\text{SPIN, Holm-Bonferroni}} = 0.02$) and receiver compression efficiency ($r = 0.14$, $p_{\text{SPIN, Holm-Bonferroni}} = 0.046$), consistent with the notion that myelin reduces conduction delay and promotes the efficient trade-off between signal rate and fidelity (reducing the transmission rate while preserving fidelity). Cortical areal scaling in neurodevelopment reflects patterns of evolutionary remodeling. Brain regions that have higher sender compression efficiency tend to disproportionately expand in relation to total brain size during neurodevelopment ($r = 0.14$, $df = 358$, $p_{\text{SPIN, Holm-Bonferroni}} = 0.045$). We observed that cortical regions with the lowest receiver compression efficiency, placing a premium on information processing fidelity, tend to disproportionately expand in relation to whole brain growth ($r = -0.20$, $df = 358$, $p_{\text{SPIN, Holm-Bonferroni}} = 0.03$). Positively scaling regions that prioritize compression-efficient broadcasting of messages arriving with high fidelity may reflect evolutionary expansion of brain regions with high information processing capacity, whereas negatively scaling regions that prioritize high-fidelity broadcasting of compressed messages may permit other modes of material, spatial, and metabolic cost efficiency.

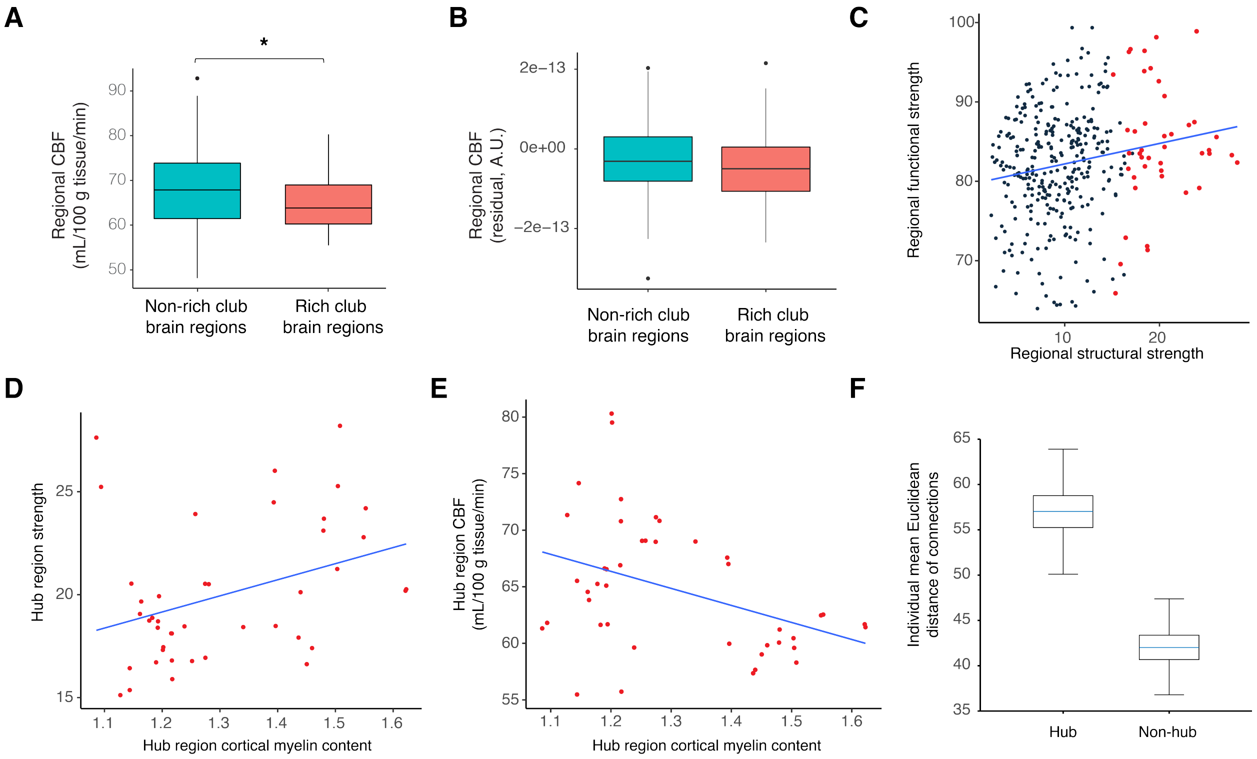


Figure 8: **No evidence of rich-club hubs incurring greater metabolic cost than other brain regions.** **(A)** We found that rich club regions tended to have reduced CBF compared to other brain regions ($W = 8420$, $p = 0.01$). Our finding that rich club regions tended to have decreased metabolic expenditure counters current understanding [66, 67, 68, 39]. However, metabolic efficiency associated with rich-club hubs is more consistent with (i) principles of evolutionary efficiency, (ii) the previously published observation that higher integrity structural connections relates to reduced metabolic expenditure (which we replicate in Main Text Figure 2), and (iii) the notion that rich-club hubs are interlinked by long-distance and myelinated connections with greater metabolic efficiency [1, 48, 49, 50, 51]. **(B)** The CBF of regions comprising the rich club was not significantly different than the CBF of other regions, when controlling for age, sex, age-by-sex interaction, and in-scanner motion ($W = 7748$, $p = 0.15$). In a sensitivity analysis only including age, sex, and age-by-sex interaction, we did not find a significant difference between the CBF of rich-club hubs and the CBF of other regions ($W = 7208$, $p = 0.54$). Given the importance of neurodevelopment to the level of both CBF and intracortical myelin in hub regions [36, 42], controlling for age and sex could explain the difference between CBF in hubs and non-hubs, if the reduced CBF is associated with increased intracortical myelin. **(C)** The hypothesis that *structural* hubs demand greater metabolic expenditure [66, 67, 68, 39] depends on taking the finding that *functional* hubs demand greater metabolic expenditure [35] and generalizing that finding to structure. Initially, this generalization might seem justified because functional connectivity explains a portion of the variance in structural connectivity [37, 38]. The lack of evidence in our data for this running hypothesis may indicate that such a generalization is not justified, particularly given heterogeneous function-structure coupling [64, 38]. Turning to our data, we found that the average structural strength is positively correlated with the functional strength (Spearman's $\rho = 0.17$, $p < 0.001$), consistent with prior reports [38]. This relationship suggests that function explains only about 2.9% of variance in structure. Hence, we may not have found evidence supporting the running hypothesis because the hypothesis relies on an imprecise assumption of a strong linear relationship between function and structure. **(D)** We next assessed whether differential cortical myelin levels could explain why the findings in Figure 8A and 8B in the Supplement both do not support the hypothesis that rich-club hubs have greater metabolic expenditure than non-hubs [39]. The hubs of the rich club (colored in red) which were stronger tended to have greater cortical myelin content (Spearman's $\rho = 0.35$, $p = 0.02$). **(E)** The hubs with greater cortical myelin content tended to have reduced metabolic expenditure (Spearman's $\rho = -0.34$, $p = 0.03$). Hence, intracortical myelin may explain why we found reduced CBF in hubs. The importance of development in the level of hub intracortical myelin may explain why we found that age and sex explain away the reduction. **(F)** Here we depict 1,041 individual structural connectomes to compare the average Euclidean distance of hub connections versus that of non-hub connections. The average hub connection has greater length than the average non-hub connection ($t = 138.7$, $df = 2080$, $p < 0.001$).

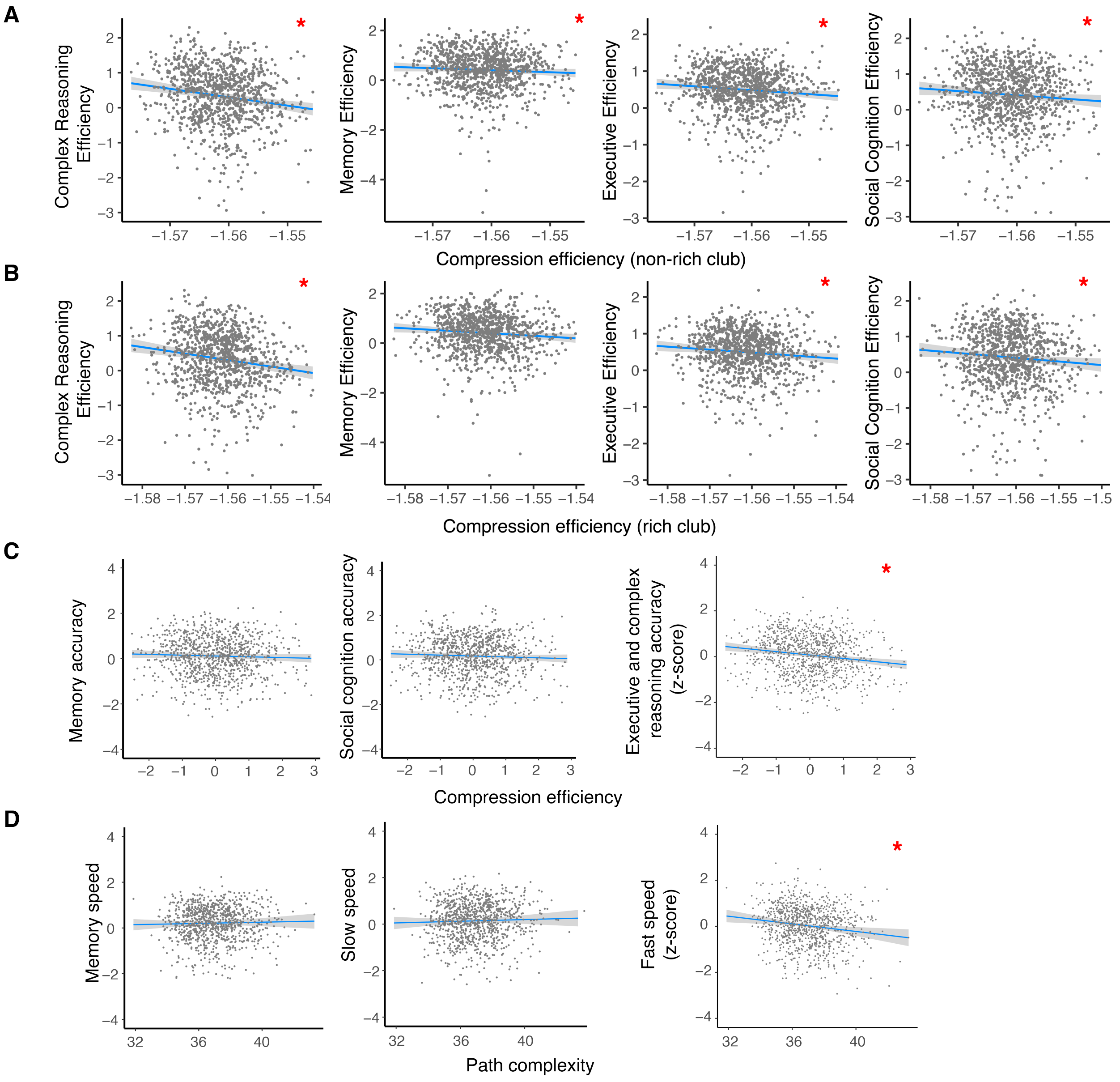


Figure 9: **Individuals with brain networks that prioritize fidelity tend to have greater cognitive efficiency. (A)** Cognitive efficiency scores depicted were residualized using generalized additive models with penalized splines controlling for global efficiency, age, sex, age-by-sex interaction, node degree, network density, and in-scanner motion. The first row depicts partial correlations with non-rich-club compression efficiency. **(B)** The second row depicts partial correlations with rich-club compression efficiency. **(C)** Network structures associated with slow-speed cognitive and memory task reaction times. Cognitive accuracy and speed scores depicted were residualized using generalized additive models with penalized splines controlling for global efficiency, age, sex, age-by-sex interaction, node degree, network density, and in-scanner motion. We did not observe a partial correlation between compression efficiency and the accuracy of memory and social cognition tasks ($p > 0.05$). **(D)** The same model was used to assess the partial correlation between path complexity and performance speed. We did not observe a partial correlation between path complexity and the speed of either memory or slow-speed tasks ($p > 0.05$). Individuals who performed more accurately on executive and complex reasoning tasks tended to have reduced compression efficiency prioritizing transmission fidelity ($t = -4.58$, bootstrap 95% CI [-6.61, -2.52], $df = 5$, $p < 0.001$, $R^2$=0.041, bootstrap 95% CI [0.021, 0.075]). As predicted by reductions in shortest path complexity supporting a low-fidelity regime of lossy compression and storage savings, individuals with quicker reaction times on tasks demanding fast speed tended to have reduced shortest path complexity (t=-2.12, bootstrap 95% CI [-4.17, -0.05], $df = 5$, $p = 0.03$, $R^2$=0.019, bootstrap 95% CI [0.01, 0.044]).

various meso-scale measures carry information about which communication process might be most efficient for a given network. For example, small world networks include “shortcuts” that reduce the number of wires needed to form the shortest paths between brain regions. Yet, they do not fully minimize wiring, implying an important functional role of the longer connections that allow the neural system to tolerate additional cost [44, 74]. Formally, small-worldness can be characterized by a statistic that combines the clustering coefficient and the global efficiency [79]. The latter is a measure of the average shortest path length, and is commonly interpreted as reflecting a network's capacity for integrated or coordinated network communication [42, 80, 81]. The former is a measure of local triangular closure, and is commonly interpreted as reflecting the network's capacity for segregated or encapsulated network communication.

In our analysis, we consider two meso-scale structures commonly observed in brain networks: modularity and the rich club. Importantly, modular brain networks also tend to have the small-world property [82]. Some modular networks – like the brain – contain a rich club, which is a set of highly connected regions representing a “backbone” of information transmission [67]. Rich-club hubs are thought to be uniquely positioned to connect modules, thereby having the marked capacity to integrate transmodal information. Those connector hubs commonly display high betweenness centrality, which is a measure of the number of different shortest paths intersecting the region that collectively support information integration.

The relationships among small-world architecture, rich-club organization, and communication have been previously considered in a descriptive manner [83, 10]. Here, we demonstrate that our model of communication moves from description to explanation and prediction. Our theory provides reasons for *why* we observe these architectural features in brain networks; those reasons stem from constraints on communication capacity and the computational principle of lossy compression. Rather than viewing shortest path and diffusion communication processes as opposing extremes on a spectrum of communication models, we synthesize both models using principles of efficient coding. The principles of efficient coding are the very same principles underlying lossy compression of information, suggesting a computational model of network integration of information.

Information transmission with random walkers is guided by a network's meso-scale structure, with properties including modularity, rich-club hubs, and path transitivity [68, 18]. Modularity measures the connectivity strength within groups of brain regions and is thought to support functional specialization by concentrating diffusion within subnetworks [84, 85, 86]. We find an exponentially increasing information processing demand that may constrain the number of brain parcels or modules. This constraint may provide a bound for accounts of the mind as massively modular [87] and design constraints on cortical parcellations typically delineating 150-200 cortical areas in each human hemisphere [88, 89, 90]. While information transfer is improved when modular circuits perform specialized functions [85], a constraint on network complexity is in line with the neural re-use hypothesis of flexible and plastic circuitry [91, 84]. In hierarchically modular brain networks [92], resource constraints could drive a compromise between the size of the network, the number of modular subnetworks, and the size of the modular subnetworks. Efficient coding in brain networks can be achieved by balancing communication fidelity and compression efficiency subject to connectome topology, plasticity, and size [88, 93, 94, 42].

Accumulating evidence supports the hypothesis that overarching principles of connectome organization underpin the spatially distributed linear dynamics of neural activity at the macro-, meso-, and micro-scales [75, 88, 95, 96, 97, 77]. In this paper, we focus on generalizing principles of neurotransmission established at the micro-scale to the macroscale using a simple linear process of random walk dynamics, consistent with the work studying the long-term behavior of linear processes subject to the structure of the connectome [18, 20, 17, 22, 13, 77]. Towards the goal of investigating micro- and macroscale biological properties of the connectome that may influence random walk dynamics, it is an exciting direction of future research to study how transmitted information is transformed by propagation along the genetic, cytoarchitectonic, and functional cortical hierarchy [78, 98]. Such studies could use similar biased random walk methods as described in Main Text Method 7.5.7.

#### 2.3 Rich-club hubs

The metabolic running costs of meso-scale structural properties, such as modularity and hubs, are not understood [92, 86, 68, 99]. Our analyses provide tools to explain how apparently materially, spatially, and metabolically expensive structural hubs are consistent with evolutionary pressures selecting for efficiency and energy minimization [100, 101, 67, 68, 102, 39]. Future studies could investigate information processing by identifying central nodes of brain networks using other methods of detecting hubs and using different measures of connection diversity, such as the participation coefficient or diverse club [64, 58].

While we replicate several findings in the literature (Supplementary Figures 4 and 5), we do not find evidence that rich-club hubs have high metabolic costs, in contrast to current understanding (Supplementary Figure 8) [66, 67]. Rather, we find weak evidence that the metabolic cost associated with rich-club hubs may be reduced compared to other brain regions during a task-unconstrained state (Supplementary Figures 8A-B) [100, 101, 1]. This null finding alleviates tension between high-cost hubs with constraints of evolutionary efficiency, suggesting that rich-club hubs may incur early material and spatial costs in negotiation for more efficient metabolic running cost. We additionally note that this finding is more consistent with reports that rich-club hubs are associated with long-distance and highly myelinated connections displaying greater metabolic efficiency [48, 1, 49, 50, 51]. Our finding is statistically robust and well-powered, using a large cohort of developing youths 20-fold larger than previous reports. Within this cohort, we consider individual differences in brain network architecture and metabolism, whereas prior analyses relied on a reference atlas [67]. Taken together, our results, prior literature, and evolutionary theory provide no evidence of a greater metabolic cost for rich-club hubs compared to other brain regions.

The efficient coding model indicated that hubs compress information, enriching the hypothesis that hubs integrate information due to their degree distribution [66, 67, 68]. Consistent with prior findings on the information efficiency of hubs [67], we found that hubs are regions prioritizing high-fidelity broadcasting of compressed input messages. Consistent with the prediction of efficient coding that neural resources allocate to the physical distribution of information, hubs receive information at reduced (compressed) rates and tend to disproportionately shrink in relation to total brain size during development. This relative shrinking is consistent with prior work suggesting that brain network development in adolescence optimizes the performance of hubs by synaptic remodeling [42]. Synaptic remodeling reduces cortical thickness and increases intracortical myelination to minimize conduction time of electrical signals or to increase the rate of information transmission [103, 42].

However, in contrast to prior hypotheses, we found that hubs were more metabolically efficient than previously thought [39, 68, 40]. Notably, our results are at odds with a small, exploratory literature that found higher metabolic cost of hubs compared to non-hubs [39]. Yet, our results are readily explained by the efficient coding hypothesis that the brain transmits maximal information in a metabolically economical and compressed form to improve future behavior [104]. Subsequent literature which found a greater enrichment of genes for metabolism in hubs can be reinterpreted in light of the lack of evidence for greater metabolic cost of rich-club hubs [39, 102, 105, 106, 107]. Prior reports of hubs exhibiting metabolically active gene expression may instead reflect the need for regulatory control of high metabolic capacity in these regions, rather than greater metabolic expenditure in general [102]. A high capacity supports efficient performance of cognitively demanding tasks which require transiently greater metabolic demand in functional connectivity hubs [108, 35]. Notably, our findings suggest that rich-club hubs may have high energy capacity but use this capacity for cost-efficient information processing. In support of high energy capacity, brain regions with greater myelin content may also contain more mitochondria from increased regional volume, axon diameter or length, or prevalence of astrocyte processes [103, 109]. Although the material and spatial cost of hubs decrease with development [42], long-run metabolic savings might depend on frequent usage of hubs to compensate for the expense of maintaining their structure and connectivity [48, 52]. Our findings are consistent with the alternative hypothesis that rich-club hubs use high energy capacity efficiently to process information.

Prior studies of rich-club connectivity have found that the organization arises from a trade-off between conserving materials and facilitating information processing [67]. The long-distance local connections of hubs have greater wiring cost but compose many shortest-paths of the brain network [67], contributing to the global efficiency of small-world architecture defined by simultaneously high clustering and low path length [110]. With increasing network size, both small-worldness and hierarchical modularity confer reduced wiring cost despite increased transitivity (clustering) [46, 110, 92]. With our macroscale efficient coding model, we reformulate interpretations of hub information integration and efficiency. In a hierarchically modular network, hubs have the least transitivity, supporting a low-fidelity regime and lossy compression. Efficient hub computation, connectivity, and consolidation may explain why the structure is heritable [83, 40].

#### 2.4 Channel Capacity

Compression efficiency quantifies the minimum rate at which information must be transmitted to reliably achieve a particular level of fidelity [111]. Every communication system also has an upper limit on the rate of reliable transmission called the channel capacity, related to information content of its input [27, 112, 63] (see Supplementary Discussion). Approaching the capacity limit is analogous to only requiring the transmission of one message for communication, as posited by models of shortest-path routing [3]. Future research could build upon our work by investigating the biological validity of error-correcting codes [27, 63, 6]. Measuring the information rate of macroscale neural dynamics may provide empirical benchmarks to test the theoretical bounds for reliable communication derived from only connectome architecture [113].

Interestingly, the channel capacity offers an alternative to the intuition that arbitrarily low probability of error is only achievable if the information rate increases indefinitely, because the channel can, in theory, transmit information with arbitrarily low error using a finite information rate. To this end, coding schemes that are more sophisticated than repetition coding have been required [27]. Determining the empirical capacity of a noisy channel will require measurements of the maximum rate at which information over all possible sources can be reliably transmitted, constrained by the physical properties of the connectome [63]. With neural representations of finite or low dimension [72, 114, 115, 116], assumptions of discrete input information makes the problem more tractable by narrowing the consideration of possible sources to a finite set of neural components or modes [117, 72, 114, 59, 118]. In addition to the complexity of the inputs or environmental stimuli [111, 73, 119], capacity is directly related to the electrophysiological properties of the connectome, including the bandwidth of oscillatory activity and the gain in neural activity [117, 9]. Measuring the information rate of macroscale neural dynamics may provide empirical benchmarks to test the theoretical predictions of the upper and lower bounds for reliable communication derived from only connectome architecture [113].

#### 2.5 Limitations and future outlook

Our model would additionally benefit from fine-tuning the distortion function. For instance, distortion could be defined by cascade duration of propagating signals [120, 98]. Moreover, fitting compression efficiency slopes can be made arbitrary by continuously adding data with distortion values approaching 0. We illustrated evidence of an upper bound using published distortion values for digital storage devices (Supplementary Figure 3D) [63]. However, the slopes are also influenced by our choice to fit a midpoint of the distribution at 50% distortion, as in prior work using the midpoint as a reference [44]. A more theoretically grounded fixed point may be 0 resources at 100% distortion, which would not change the interpretation of compression efficiency as quantifying the prioritization of fidelity or compression, but may affect the interpretation of cost-dependent errors. Linking theory with measurements, such as recent work describing transformations of neural information along structural pathways [113], is a promising direction of research.
